# Streamlined use of protein structures in variant analysis

**DOI:** 10.1101/2021.09.10.459756

**Authors:** Sandeep Kaur, Neblina Sikta, Andrea Schafferhans, Nicola Bordin, Mark J. Cowley, David M. Thomas, Mandy L. Ballinger, Seán I. O’Donoghue

**Affiliations:** Garvan Institute of Medical Research, Sydney, Australia; University of New South Wales, Sydney, Australia; Department of Bioengineering Sciences, Weihenstephan-Tr. University of Applied Sciences, Freising, Germany; Department of Informatics, Bioinformatics & Computational Biology, Technical University of Munich, Germany; Institute of Structural and Molecular Biology, University College London, UK; Children’s Cancer Institute, Sydney, NSW, Australia; CSIRO Data61, Sydney, Australia

**Keywords:** Variant analysis, Data integration, Protein structures, Molecular graphics

## Abstract

**Motivation:** Variant analysis is a core task in bioinformatics that requires integrating data from many sources. This process can be helped by using 3D structures of proteins, which can provide a spatial context that can provide insight into how variants affect function. Many available tools can help with mapping variants onto structures; but each has specific restrictions, with the result that many researchers fail to benefit from valuable insights that could be gained from structural data.

**Results:** To address this, we have created a streamlined system for incorporating 3D structures into variant analysis. Variants can be easily specified via URLs that are easily readable and writable, and use the notation recommended by the Human Genome Variation Society (HGVS). For example, ‘https://aquaria.app/SARS-CoV-2/S/?N501Y’ specifies the N501Y variant of SARS-CoV-2 S protein. In addition to mapping variants onto structures, our system provides summary information from multiple external resources, including COSMIC, CATH-FunVar, and PredictProtein. Furthermore, our system identifies and summarizes structures containing the variant, as well as the variant-position. Our system supports essentially any mutation for any well-studied protein, and uses all available structural data — including models inferred via very remote homology — integrated into a system that is fast and simple to use. By giving researchers easy, streamlined access to a wealth of structural information during variant analysis, our system will help in revealing novel insights into the molecular mechanisms underlying protein function in health and disease.

**Availability:** Our resource is freely available at the project home page (https://aquaria.app). After peer review, the code will be openly available via a GPL version 2 license at https://github.com/ODonoghueLab/Aquaria. PSSH2, the database of sequence-to-structure alignments, is also freely available for download at https://zenodo.org/record/4279164.

**Supplementary information:** None.

## Introduction

For many clinicians and researchers, one of their core tasks involves understanding the impact of a variant on a protein’s function. Gaining this understanding can be a complex process, requiring integration of data from clinical reports, *in vitro* experiments, and *in silico* predictions. In many cases, this integration can be helped by using 3D structures of proteins, which can provide detailed insight into functional impacts of variants — especially how interactions with other molecules may be affected (Gress 2020). One of the main methods for using these structures is simply as a location to map variant data; this then allows a researcher to consider spatial context when assessing functional impact.

Many tools are available to help researchers derive insights from 3D structure (Glusman et al. 2017). Some provide a streamlined workflow, but are mostly restricted to human proteins (e.g., MuPIT Interactive from Niknafs et al. 2013; VarSite from Laskowski et al. 2020); others are restricted to fixed sets of variants (e.g., MISCAST from Iqbal et al. 2020); many tools (e.g., dSysMap from Mosca et al. 2015; Mechismo from Betts et al. 2015) are mostly restricted to using only experimentally derived 3D structures (i.e., the PDB; Berman et al. 2000). Other tools provide rich, generic capabilities with few restrictions, but can be complex and time-consuming to use (e.g., ChimeraX from Pettersen et al. 2020).

Unfortunately, as a result of these limitations, structural data are often not included in variant analysis, and often not seen as essential in variant analysis (e.g., Pabinger et al. 2014). This is especially true in clinical genomics, where researchers are often under time pressure to analyze large numbers of variants. Thus, unfortunately, many researchers fail to benefit from the potentially valuable insights that could be gained from structural data.

Addressing this issue was our core goal in this work, which arose from a collaboration between bioinformaticians and clinical genomics researchers. As a first step, we realized the need to devise a very simple entry point for including structural data in variant analysis; this entry point needs to be easily added to existing genomics workflows — much of which involves spreadsheets, as a major format for management and communication of mutational data.

In addition, we wanted to overcome limitations with existing tools, as mentioned above; thus, we aimed to create a system that supports essentially any mutation for any protein, and lets researchers use all available structural data — including models inferred via very remote homology. Finally, our system should be easy to learn, and be fast and simple to use.

To address these goals, we built upon Aquaria, a web-based, molecular graphics resource (O’Donoghue et al. 2015) that is freely available and open-source. Aquaria provides structural information for >500,000 protein sequences in SwissProt (The UniProt Consortium 2019), which includes proteins from many model organisms used in research. For each protein sequence, Aquaria uses HHblits (Steinegger et al. 2019) to pre-calculate alignments onto all available 3D structures. This generates >120 million sequence-to-structure alignments (Schafferhans and O’Donoghue 2020), which gives an average of ∼200 structural models for each protein sequence.

In this work, we describe extensions to Aquaria that largely address the above limitations with existing structure-based variant analysis tools. The extensions include summarizing of structures containing the variant, incorporating new variant data from COSMIC (Tate et al. 2019) and FunVar (Sillitoe et al. 2021), as well as improving how data are integrated and presented to the user — including data from UniProt (The UniProt Consortium 2019), SNAP2 (Hecht, Bromberg, and Rost 2015), and PredictProtein (Yachdav et al. 2014). We demonstrate these capabilities by analysing variants in two case studies: the human oncoprotein NRAS and the S protein from SARS-CoV-2.

## Methods

### Variant notation

For specifying variants, our system uses the <u>HGVS notation</u> (den Dunnen et al. 2016), which is widely used in clinical genomics. Our system supports essentially all variants that can be expressed using the notation (**Table 1**); the only exceptions are cases where position is not specified — for example ‘p.0’, which indicates that no protein is produced, or ‘[?]’, which indicates an allele with unspecified consequence — or cases where the mutation is silent (e.g. ‘p.Cys370=’). HGVS notation was developed for annotating genomics data; to increase convenience with protein sequences, we added two optional rules as minor extensions: (1) the ‘p.’ prefix optional, and (2) amino acids can be abbreviated using either three letter or single letter coding. Thus, in our system, ‘C370T’ is equivalent to ‘p.Cys370Thr’. Currently, our system does not support annotation of DNA or RNA 3D structures; however, we are plan to support this later, in which case, mutations on such sequences will require the prefixes “c.”, “g.”, “m.”, “n.”, “o.”, or “r.” to specify numbering for coding DNA, genomes, mitochondrial DNA, non-coding DNA, circular genomes, or for RNA, respectively.

**Table 1.**
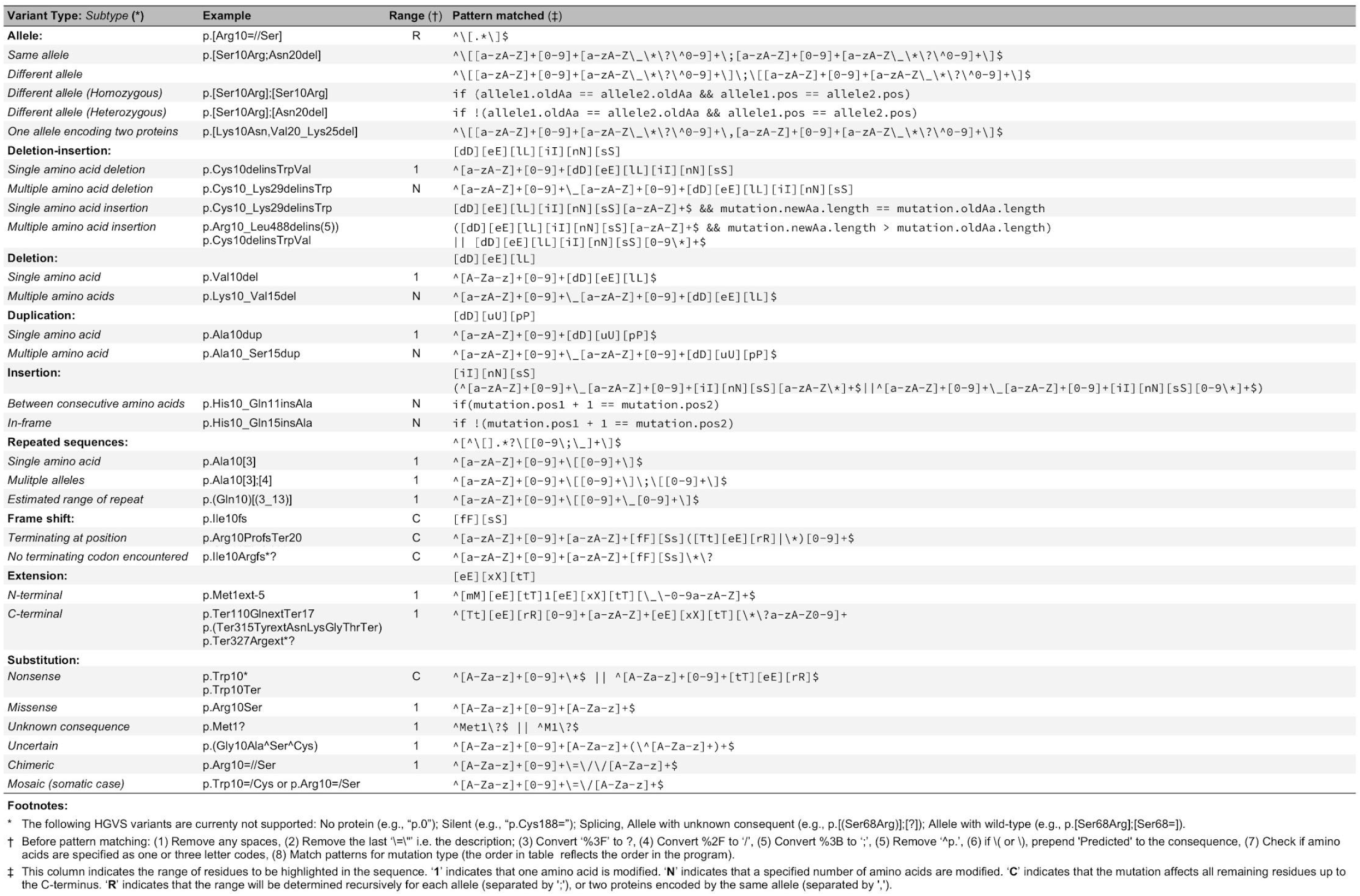
Notation for variants currently supported by Aquaria’s URL schema

### Architecture

Aquaria (O’Donoghue et al. 2015) is built mainly using JavaScript; the server uses Node.js, while the client uses primarily Vue.js and Jolecule (https://jolecule.com). Underlying Aquaria is PSSH2, a Sequel database of sequence-to-structure alignments (Schafferhans and O’Donoghue 2020). We have recently completed a major refactoring of the Aquaria user interface, which we plan to describe in a forthcoming publication. Below, we describe the new variant analysis capabilities. When a variant is specified by the user in the URL, Aquaria first identifies the consequence according to the patterns described in **Table 1**. In parallel, Aquaria fetches feature sets from several external sources (below and **Figure 1**); once data from a source is received, it is extracted and converted to Aquaria’s feature format (https://bit.ly/aquaria-features), and the queried variants are then checked to see if any of the features pertain to it. The features containing the variant are added to a new ‘Added features’ set, which is updated every time data from an external source is received. Upon initial page load, the ‘Added features’ are displayed on the 3D structure; however, the user can manually switch to any other feature set. It is worth noting that the mutation is specified via the URL’s query string (i.e., following the ‘?’ character), which gets sent to the Aquaria server; however, all feature classification, requests to external servers, and processing are performed on the user’s browser (client-side). On the server side, the query is used to identify the best matching structure for initial display in Jolecule (the 3D viewer). Additionally, structure counts pertaining to the specified variant are determined (described in the next section).

**Fig. 1.**
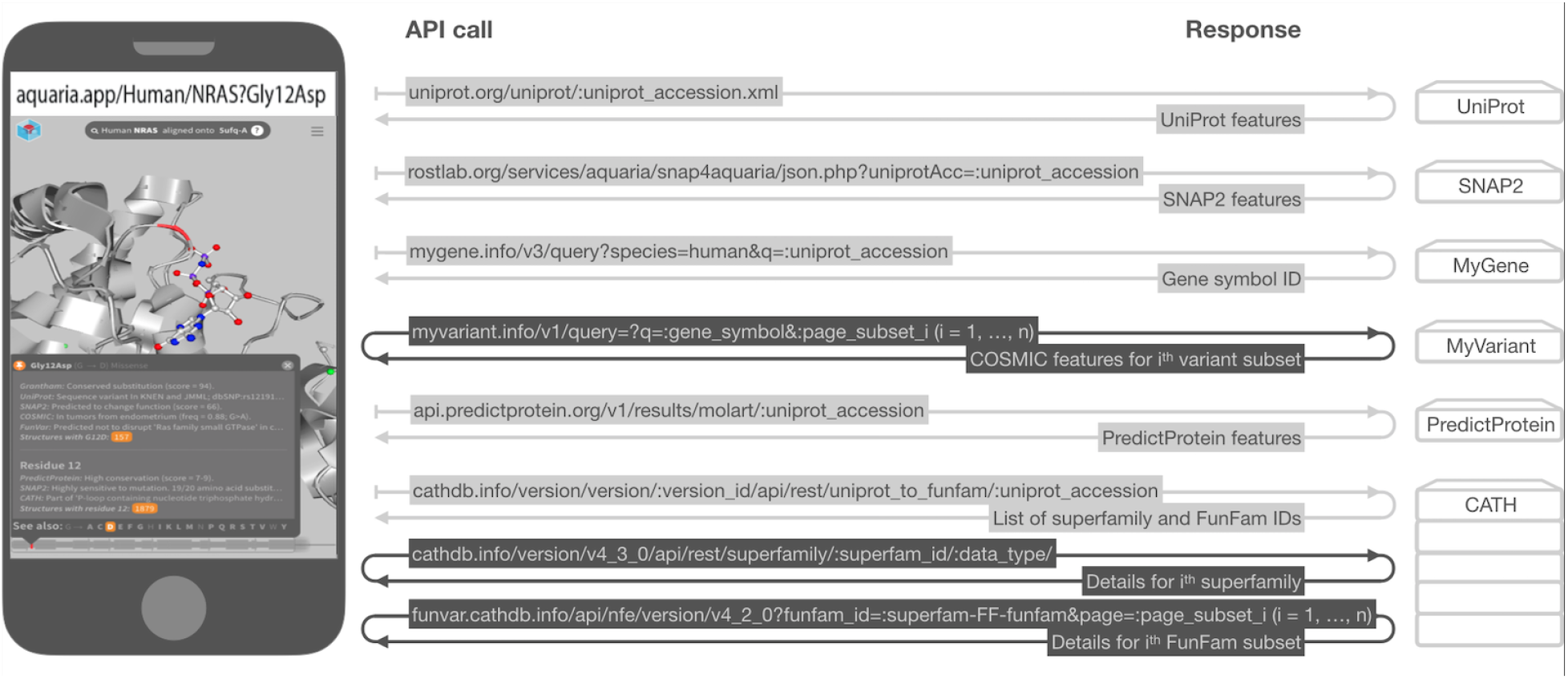
API calls to external servers used to retrieve variant information shown in the popup on an Aquaria client. API calls shown with light grey coloring are called only once. Dark coloring indicates repeated API calls, which are used to iterate through lists of features.

### Determining structure counts and choosing the best structure

On the server, the query is used to identify the best matching structure most relevant to the specified variants, from the list of all matching structures for a sequence (which were identified via homology modelling (Schafferhans and O’Donoghue 2020)) for initial display in Jolecule, as follows:

1. If a user specifies a PDB identifier in the URL, then the specified PDB is chosen for display.
2. Otherwise, when multiple variants are specified, a subset of structures is identified which covers the most number of variants, starting from the left-most user-specified (i.e. in the order specified by the user in the URL).
3. If any specified variants are ‘missense’ variants, then the subset is filtered further to identify structures containing the new amino-acid resulting from the missense variant.
4. If multiple structures result from the previous two filtering steps, then the structure with the highest rank, based on alignment identity score, coverage score, resolution and experimental method, is chosen from the filtered list.
5. If no structure results from (2) and (3), then the default best matching structure is added (i.e. the highest ranked matching structure for the sequence).

For each specified missense variant the number of structures containing the new amino acid resulting from the mutation are identified. Additionally, the number of structures for a variant’s sequence position are also identified and returned to the client. These are then added to the ‘Added features’ feature set, and are available for visualization and navigation via the popup.

### UniProt

The UniProt Knowledgebase (The UniProt Consortium 2019), consists of, perhaps the largest collection of protein sequences and their published curated annotations that describe regions or sites of interest in the protein sequence such as secondary structure regions or PTM sites. Aquaria fetches and displays the available curated feature sets via UniProt’s API (**Figure 1**). If the information for the user-specified variant in the URL is available in the following UniProt feature sets, namely ‘Modified residue’ (modified residues excluding lipids, glycans and protein cross-links), ‘Mutagenesis site’ (site which has been experimentally altered by mutagenesis), ‘Sequence conflict’ (description of sequence discrepancies of unknown origin), ‘Sequence variant’ (description of a natural variant of the protein), ‘Cross-link’ (residues participating in covalent linkage(s) between proteins), ‘Site’ (interesting single amino acid site on the sequence) or ‘Metal ion-binding site’ (binding site for a metal ion); these data are then collected and added to the ‘Added features’ feature set.

#### COSMIC

The Catalogue of Somatic Mutations in Cancer (COSMIC) (Tate et al. 2019) is a curated resource of somatic human cancers. The data are collected from literature curation of genes and genome-wide screens. To access these data, Aquaria fetches a list of all variants for a given protein available in COSMIC v68 (1,024,494 total variants) from MyVariant (**Figure 1**). In the returned data, the variants are described at the genome level only; thus, Aquaria then uses data from SnpEff (Cingolani et al. 2012) to extract the corresponding HGVS protein description and variation effect, which determine the sequence highlighting for these variants, e.g. for ‘start lost’ mutations, the entire sequence is highlighted indicating that the entire protein is affected. If a COSMIC variant exists at the same position as the user-specified variant in the URL, then these data are collected and added to the ‘Added features’ feature set.

#### SNAP2

SNAP2 (Hecht, Bromberg, and Rost 2015) is an *in silico* prediction method that uses neural networks to predict effects of single amino acid substitutions on protein functions. For each residue in a protein sequence, SNAP2 provides two types of scores. The first, mutational sensitivity, consists of a list of 20 scores that indicate the predicted functional consequences of the position being occupied by each of the 20 standard amino acids. Large, positive scores (up to 100) indicate substitutions likely to have deleterious changes, while negative scores (down to -100) indicate no likely functional change. A single summary score is then calculated based on the total fraction of substitutions predicted to have deleterious effect, taken to be those with a score > 40. The second, mutational score, is based on the same 20 scores above, but calculates a single summary score for each residue as the average of the individual scores for each of the 20 standard amino acids. Aquaria fetches and displays these scores via the SNAP2 API (**Figure 1**). For a user-specified variant, Aquaria adds the variant-specific SNAP2 prediction as well as the average mutational sensitivity for that position to the ‘Added features’ feature set.

#### CATH

The CATH resource (Sillitoe et al. 2021) identifies and classifies structural and functional domains in proteins using a hierarchical classification that consists of class (secondary structure content), architecture (orientation of secondary structure), topology (shape and connectivity of secondary structures), homologous superfamily and functional family (functionally coherent groups). Aquaria fetches CATH annotations for a protein using multiple API-endpoints (https://github.com/UCLOrengoGroup/cath-api-docs) and makes them available as two feature sets namely, superfamilies and functional families, both of which contain the class, architecture and topology classification, along with gene ontology, enzyme commission terms and species distributions of the homologous superfamily. The functional family feature set additionally contains the functional family domain. Since the CATH annotations for SARS-COV-2 are not yet available via the API, we host and utilize a pre-release version of these annotations (**Figure 1**). For a user-specified variant, if the variant position forms part of any domain, then they are added to the ‘Added features’ feature set.

### FunVar

FunVar is a new resource derived from CATH and other inputs (Sillitoe et al. 2021) that presents the predicted impact of variants (FunVar), on protein structures and functions, derived by identifying whether the variants lie within or in close proximity to the interface regions or the functional sites, which are highly conserved residues in CATH functional family multiple sequence alignments. FunVar is currently available for proteins implicated in human cancers (data from (The Cancer Genome Atlas Research Network et al. 2013)) and, SARS-COV-2 viral and its human host interactor proteins (data from (Gordon et al. 2020)). Aquaria fetches and displays the FunVars using the FunVar API **(Figure 1**). If any FunVar variant exists at the same position as the user-specified variant in the URL, then these data are collected and added to the ‘Added features’ feature set.

### PredictProtein

The PredictProtein resource (Yachdav et al. 2014) provides comprehensive computational prediction regarding multiple aspects of protein structure and function. PredictProtein features include secondary structure, solvent accessibility, conservation and, protein and DNA binding. Aquaria fetches and displays these features using the PredictProtein API (**Figure 1**). For a variant specified by the user in the URL, PredictProtein’s conservation prediction and score are added to the ‘Added features’ feature set.

## Results

### Specifying variants

Our system allows variants to be specified via the following URL schema:

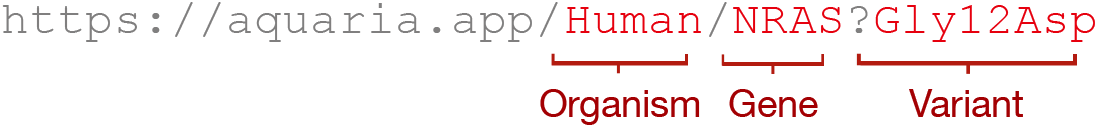

Our system supports most variants that can be described using HGVS notation (den Dunnen et al. 2016), with minor exceptions as noted in **Table 1**. The URL schema can also be used to specify a description for each variant, as shown below. The description can include HTML tags, thus allowing inclusion of images or interactive graphics.

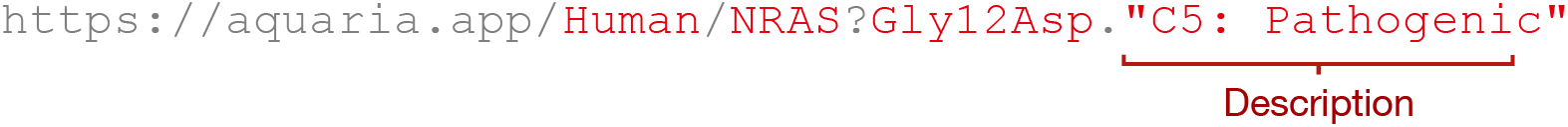

Finally, the schema can also be used to specify multiple variants for one protein:

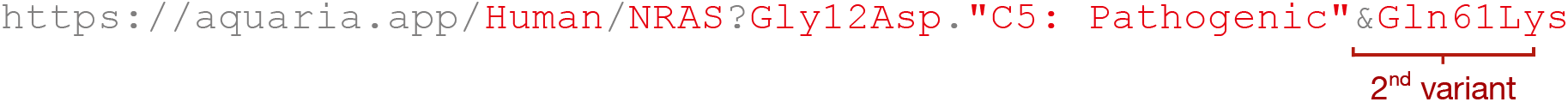

The systematic pattern of the URL enables it to be easily used within spreadsheets - a common tool in clinical genomics - containing mutational data from patients. The spreadsheet formula below illustrates how this can be done by referencing cells that contain the gene name and the variant (in this example, cells A2 and B2, respectively):

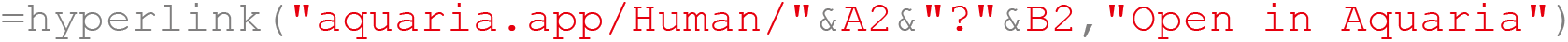

Clicking on a link that follows the above URL schema opens a web page on the Aquaria resource that gives access to all 3D structures related to the specified protein, and shows the specified protein mapped both, onto a representation of the protein sequence, as well as the ‘best’ matching structure. See next section for details.

### Exploring variants in 3D

Clicking on a link matching the above URL schema opens the Aquaria.app web resource focused on the specified protein, and showing the spatial context, sequence position, and supporting information for the specified variant (**Figure 2**). Each protein has on average ∼200 structural models; Aquaria automatically selects and displays a ‘best’ 3D structure (O’Donoghue et al. 2015). Each specified variant is indicated using red highlighting in both the structural coverage map (**Figure 2B** & **E**) and on the structure (**Figure 2A**). The structural coverage map is assembled by overlaying the sequence maps of all the top matching structures. It summarises and indicates all the structural information that is present for the sequence, i.e. what is known or not known about the 3D structure of the protein sequence. The highlighted residues are annotated with information automatically fetched from external sources, including UniProt (The UniProt Consortium 2019), COSMIC (Tate et al. 2019), SNAP2 (Hecht, Bromberg, and Rost 2015), PredictProtein (Yachdav et al. 2014) and FunVar (Sillitoe et al. 2021) (e.g. **Figure 2C and D**). This information is revealed on hover in a popup (**Figure 2** and **Figure 3**).

**Fig. 2.**
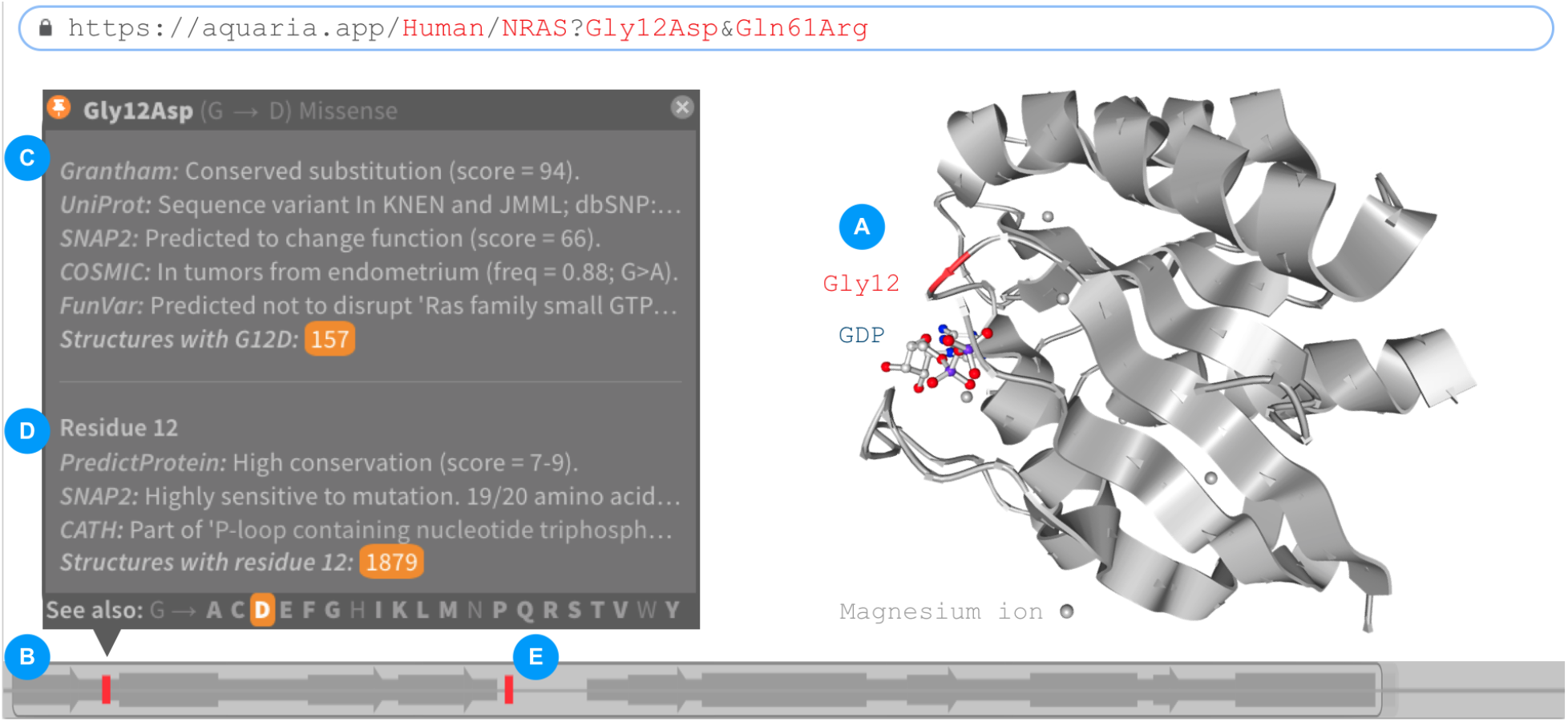
Aquaria view of two cancer-driving variants in human NRAS. (**A**) Residue Gly12 is close to a GTP binding site; mutations in this residue may affect this critical interaction. (**B**) The sequence position of each variant is shown in red; hovering over a variant opens a popup. (**C**) Popup showing variant information aggregated from multiple sources. (**D**) Shows information about the residue position. (**E**) One of the specified variants (Gln61Arg) occurs in a region that is missing in this PDB entry (3con; Nedyalkova et al. 2017). Made using Aquaria and edited in Keynote.

**Fig. 3.**
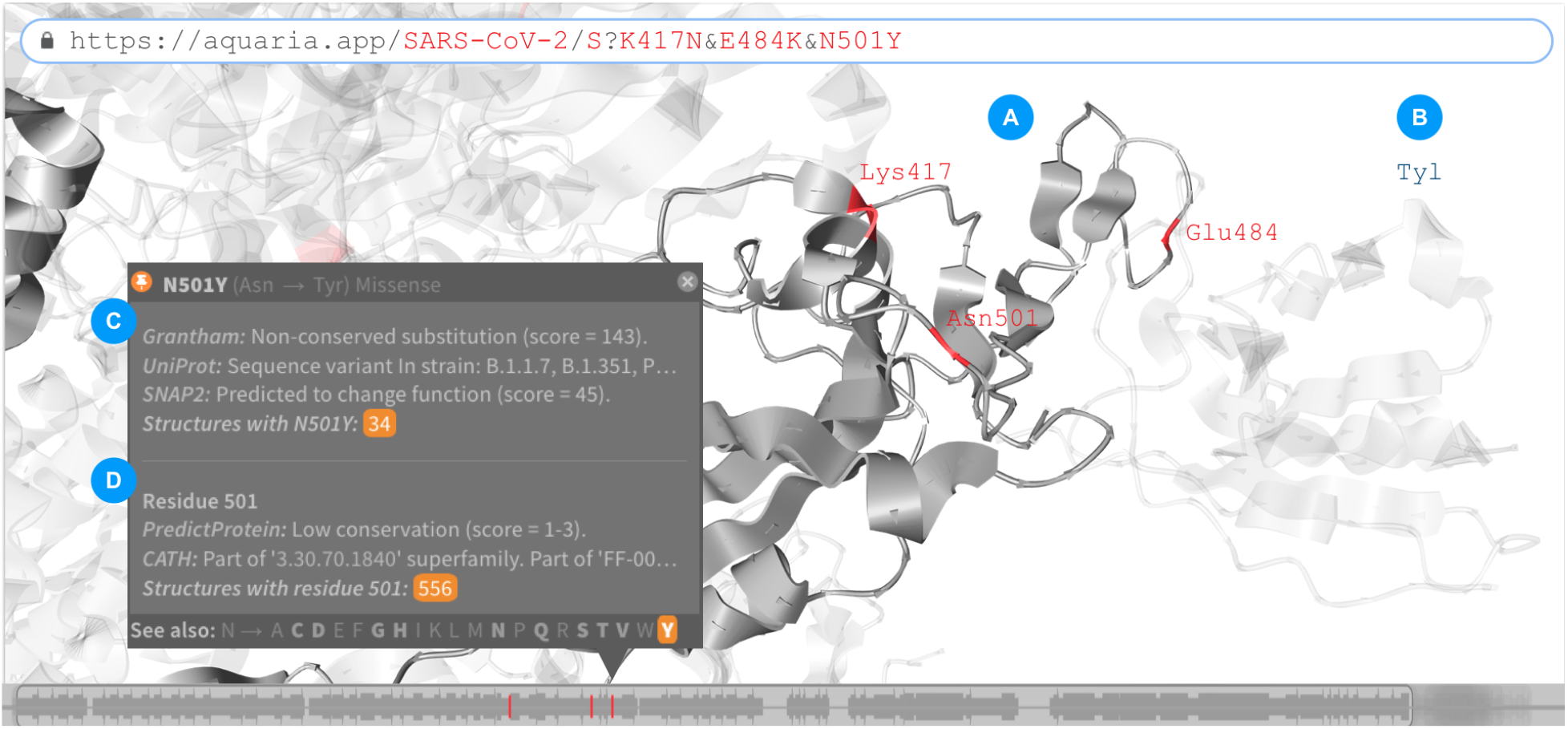
Aquaria view of three variants in SARS-CoV-2 S protein. (**A**) In this PDB entry (6zxn; Hanke et al. 2020), only one of the variants (E484K) occurs very closely to a site binding with ‘Tyl’ (**B**), a nanobody, shown with semi-transparent coloring, known to block binding to ACE2. Thus, the other two mutations (K417N and N501Y) may have little effect on Tyl binding. (**C**) Two sources suggest that N501Y will change function. 34 structures are available containing the variant, and the variant has been identified in a number of lineages. (**D**) This residue position has medium sensitivity to mutation, low conservation, and is part of a region involved in homooligomerization and in binding to cell surface receptors. 556 structures are available for this position. Made using Aquaria and edited in Keynote.

The 3D orientation, placement and zoom of the 3D structure can be easily manipulated using mouse controls. The initially displayed structure may not always be the most relevant to study the molecular mechanisms of interest. For example, it may not span a variant of interest. Aquaria, however, provides easy access to all 3D structures available related to any variant, variant-position and protein. To see all available structures related to a given variant or variant-position, clicking on the structure counts in the variant popup (e.g. **Figure 2** and **Figure 3**) reveals an interactive tree which displays structures containing the variant, or the variant-position. From the tree, any structure can be clicked, which choses and loads the structure - thus enabling navigation to any structure relevant to the variant. To see all available structures related to a given protein, a researcher just needs to tap or click on the icon at the bottom and centre of the screen. Aquaria will then display a concise visual overview of all matching structures; this overview helps researchers find and navigate to structures more relevant to their work (e.g finding interacting structures where the variant may lie on the interacting domain).

### Case study 1: Human NRAS

An illustration for how insight can be gained from mapped variants is provided in **Figure 2**, which shows a 3D structure of the human NRAS oncoprotein, mapped with two cancer-driving mutations (Gly12Asp and Gln61Arg). The Gly12Asp mutation is in close proximity to a bound GTP molecule (**Figure 2A**), suggesting that this mutation may interfere with GTP binding — which is critical for NRAS to function normally (Kapoor and Travesset 2015). This structure-based insight is confirmed to be correct via information shown in the popup, thus providing a validation that the information presented can be effective in assessing variants.

Aquaria also shows that the Gln61Arg mutation occurs in a region that is missing in the PDB entry shown by default; however, by using the Aquaria interface, a researcher can find other related structures where this region is present, demonstrating the value of making all related structures easily accessible. An example is provided in the URL below:

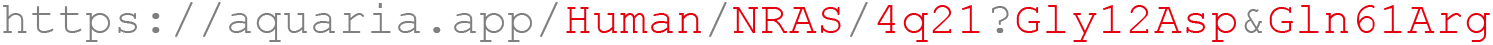

### Case study 2: SARS-CoV-2 S protein

Here, we showcase how Aquaria’s variant analysis capabilities can help in accessing three SARS-CoV-2 mutations of particular concern in the current COVID-19 pandemic (Leung et al. 2021; Tang et al. 2021; Greaney et al. 2021); the mutations can be accessed via the URL:

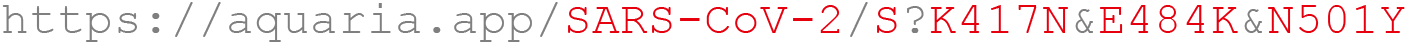

Only one of these mutations (E484K) occurs very close to a bound protein called ‘Tyl’, a nanobody known to block binding to ACE2, present in the structure shown (**Figure 3A** & **B**). Thus, we infer that E484K — but not K417N or N501Y — may block binding of this nanobody.

## Discussion

The URL schema for variant specification developed in this work is both human-readable and human-writeable; in addition to its utility in spreadsheets - enabled via standardized Aquaria URL schema and HGVS nomenclature - this improved readability compared with other related resources may help more researchers access and use structural information more routinely. To our knowledge, no other system supports essentially any mutation for any well-studied protein, uses all available structural data — including models inferred via very remote homology, summarizes structures available for a variant — integrated into a system that is fast and simple to use. By giving researchers easy, streamlined access to a wealth of structural information during variant analysis, our system will help in revealing novel insights into the molecular mechanisms underlying protein function in health and disease.

## Acknowledgements

We gratefully acknowledge the help of Paul Ashford and other members of the CATH team in advising us on accessing and interpreting FunFam and FunVar information. Also, we thank Bosco Ho for developing and maintaining Jolecule.

## References

Berman, Helen M., John Westbrook, Zukang Feng, Gary Gilliland, T. N. Bhat, Helge Weissig, Ilya N. Shindyalov, and Philip E. Bourne. 2000. “The Protein Data Bank.” Nucleic Acids Research 28 (1): 235–42. https://doi.org/10.1093/nar/28.1.235.

Betts, Matthew J., Qianhao Lu, YingYing Jiang, Armin Drusko, Oliver Wichmann, Mathias Utz, Ilse A. Valtierra-Gutiérrez, et al. 2015. “Mechismo: Predicting the Mechanistic Impact of Mutations and Modifications on Molecular Interactions.” Nucleic Acids Research 43 (2): e10. https://doi.org/10.1093/nar/gku1094.

Cingolani, Pablo, Adrian Platts, Le Lily Wang, Melissa Coon, Tung Nguyen, Luan Wang, Susan J. Land, Xiangyi Lu, and Douglas M. Ruden. 2012. “A Program for Annotating and Predicting the Effects of Single Nucleotide Polymorphisms, SnpEff: SNPs in the Genome of Drosophila Melanogaster Strain W1118; Iso-2; Iso-3.” Fly 6 (2): 80–92. https://doi.org/10.4161/fly.19695.

Dunnen Johan, T. den, Raymond Dalgleish, Donna R. Maglott, Reece K. Hart, Marc S. Greenblatt, Jean McGowan-Jordan, Anne-Francoise Roux, et al. 2016. “HGVS Recommendations for the Description of Sequence Variants: 2016 Update.” Human Mutation 37 (6): 564–69. https://doi.org/10.1002/humu.22981.

Glusman, Gustavo, Peter W. Rose, Andreas Prlić, Jennifer Dougherty, José M. Duarte, Andrew S. Hoffman, Geoffrey J. Barton, et al. 2017. “Mapping Genetic Variations to Three-Dimensional Protein Structures to Enhance Variant Interpretation: A Proposed Framework.” Genome Medicine 9 (1): 113. https://doi.org/10.1186/s13073-017-0509-y.

Gordon, David E., Gwendolyn M. Jang, Mehdi Bouhaddou, Jiewei Xu, Kirsten Obernier, Kris M. White, Matthew J. O’Meara, et al. 2020. “A SARS-CoV-2 Protein Interaction Map Reveals Targets for Drug Repurposing.” Nature 583 (7816): 459–68. https://doi.org/10.1038/s41586-020-2286-9.

Greaney, Allison J., Andrea N. Loes, Katharine H. D. Crawford, Tyler N. Starr, Keara D. Malone, Helen Y. Chu, and Jesse D. Bloom. 2021. “Comprehensive Mapping of Mutations to the SARS-CoV-2 Receptor-Binding Domain That Affect Recognition by Polyclonal Human Serum Antibodies.” BioRxiv, January, 2020.12.31.425021. https://doi.org/10.1101/2020.12.31.425021.

Gress, Alexander. 2020. “Integration of Protein Three-Dimensional Structure into the Workflow of Interpretation of Genetic Variants.” https://doi.org/10.22028/D291-32073.

Hanke, Leo, Laura Vidakovics Perez, Daniel J. Sheward, Hrishikesh Das, Tim Schulte, Ainhoa Moliner-Morro, Martin Corcoran, et al. 2020. “An Alpaca Nanobody Neutralizes SARS-CoV-2 by Blocking Receptor Interaction.” Nature Communications 11 (1): 4420. https://doi.org/10.1038/s41467-020-18174-5.

Hecht, Maximilian, Yana Bromberg, and Burkhard Rost. 2015. “Better Prediction of Functional Effects for Sequence Variants.” BMC Genomics 16 (8): S1. https://doi.org/10.1186/1471-2164-16-S8-S1.

Iqbal, Sumaiya, David Hoksza, Eduardo Pérez-Palma, Patrick May, Jakob B Jespersen, Shehab S Ahmed, Zaara T Rifat, et al. 2020. “MISCAST: MIssense Variant to Protein StruCture Analysis Web SuiTe.” Nucleic Acids Research 48 (W1): W132–39. https://doi.org/10.1093/nar/gkaa361.

Kapoor, Abhijeet, and Alex Travesset. 2015. “Differential Dynamics of RAS Isoforms in GDP- and GTP-Bound States: Differential Dynamics of RAS Isoforms.” Proteins: Structure, Function, and Bioinformatics 83 (6): 1091–1106. https://doi.org/10.1002/prot.24805.

Laskowski, Roman A., James D. Stephenson, Ian Sillitoe, Christine A. Orengo, and Janet M. Thornton. 2020. “VarSite: Disease Variants and Protein Structure.” Protein Science 29 (1): 111–19. https://doi.org/10.1002/pro.3746.

Leung, Kathy, Marcus HH Shum, Gabriel M Leung, Tommy TY Lam, and Joseph T Wu. 2021. “Early Transmissibility Assessment of the N501Y Mutant Strains of SARS-CoV-2 in the United Kingdom, October to November 2020.” Eurosurveillance 26 (1). https://doi.org/10.2807/1560-7917.ES.2020.26.1.2002106.

Mosca, Roberto, Jofre Tenorio-Laranga, Roger Olivella, Victor Alcalde, Arnaud Céol, Montserrat Soler-López, and Patrick Aloy. 2015. “DSysMap: Exploring the Edgetic Role of Disease Mutations.” Nature Methods 12 (3): 167–68. https://doi.org/10.1038/nmeth.3289.

Nedyalkova, L., Y. Tong, W. Tempel, L. Shen, P. Loppnau, C.H. Arrowsmith, A.M. Edwards, et al. 2017. “Crystal Structure of the Human NRAS GTPase Bound with GDP.” PDB unpublished raw dataset. https://dx.doi.org/10.2210/pdb3con/pdb.

Niknafs, Noushin, Dewey Kim, RyangGuk Kim, Mark Diekhans, Michael Ryan, Peter D. Stenson, David N. Cooper, and Rachel Karchin. 2013. “MuPIT Interactive: Webserver for Mapping Variant Positions to Annotated, Interactive 3D Structures.” Human Genetics 132 (11): 1235–43. https://doi.org/10.1007/s00439-013-1325-0.

O’Donoghue Seán I., Kenneth S. Sabir, Maria Kalemanov, Christian Stolte, Benjamin Wellmann, Vivian Ho, Manfred Roos, et al. 2015. “Aquaria: Simplifying Discovery and Insight from Protein Structures.” Nature Methods 12 (2): 98–99. https://doi.org/10.1038/nmeth.3258.

Pabinger, S., A. Dander, M. Fischer, R. Snajder, M. Sperk, M. Efremova, B. Krabichler, M. R. Speicher, J. Zschocke, and Z. Trajanoski. 2014. “A Survey of Tools for Variant Analysis of Next-Generation Genome Sequencing Data.” Briefings in Bioinformatics 15 (2): 256–78. https://doi.org/10.1093/bib/bbs086.

Pettersen, Eric F., Thomas D. Goddard, Conrad C. Huang, Elaine C. Meng, Gregory S. Couch, Tristan I. Croll, John H. Morris, and Thomas E. Ferrin. 2020. “CSF ChimeraX: Structure Visualization for Researchers, Educators, and Developers.” Protein Science, October, pro.3943. https://doi.org/10.1002/pro.3943.

Schafferhans, Andrea, and Sean O’Donoghue. 2020. “PSSH2 - Database of Protein Sequence-to-Structure Homologies.” Zenodo. https://doi.org/10.5281/ZENODO.4279164.

Sillitoe, Ian, Nicola Bordin, Natalie Dawson, Vaishali P Waman, Paul Ashford, Harry M Scholes, Camilla S M Pang, et al. 2021. “CATH: Increased Structural Coverage of Functional Space.” Nucleic Acids Research 49 (D1): D266–73. https://doi.org/10.1093/nar/gkaa1079.

Steinegger, Martin, Markus Meier, Milot Mirdita, Harald Vöhringer, Stephan J. Haunsberger, and Johannes Söding. 2019. “HH-Suite3 for Fast Remote Homology Detection and Deep Protein Annotation.” BMC Bioinformatics 20 (1): 473. https://doi.org/10.1186/s12859-019-3019-7.

Tang, Julian W., Oliver T.R. Toovey, Kirsty N. Harvey, and David D.S. Hui. 2021. “Introduction of the South African SARS-CoV-2 Variant 501Y.V2 into the UK.” Journal of Infection, January, S016344532100030X. https://doi.org/10.1016/j.jinf.2021.01.007.

Tate, John G, Sally Bamford, Harry C Jubb, Zbyslaw Sondka, David M Beare, Nidhi Bindal, Harry Boutselakis, et al. 2019. “COSMIC: The Catalogue Of Somatic Mutations In Cancer.” Nucleic Acids Research 47 (D1): D941–47. https://doi.org/10.1093/nar/gky1015.

The Cancer Genome Atlas Research Network, John N Weinstein, Eric A Collisson, Gordon B Mills, Kenna R Mills Shaw, Brad A Ozenberger, Kyle Ellrott, Ilya Shmulevich, Chris Sander, and Joshua M Stuart. 2013. “The Cancer Genome Atlas Pan-Cancer Analysis Project.” Nature Genetics 45 (10): 1113–20. https://doi.org/10.1038/ng.2764.

The UniProt Consortium. 2019. “UniProt: A Worldwide Hub of Protein Knowledge.” Nucleic Acids Research 47 (D1): D506–15. https://doi.org/10.1093/nar/gky1049.

Yachdav, Guy, Edda Kloppmann, Laszlo Kajan, Maximilian Hecht, Tatyana Goldberg, Tobias Hamp, Peter Hönigschmid, et al. 2014. “PredictProtein—an Open Resource for Online Prediction of Protein Structural and Functional Features.” Nucleic Acids Research 42 (W1): W337–43. https://doi.org/10.1093/nar/gku366.

